# Intolerance to uncertainty modulates neural synchrony between political partisans

**DOI:** 10.1101/2020.10.28.358051

**Authors:** Jeroen M. van Baar, David J. Halpern, Oriel FeldmanHall

## Abstract

Political partisans see the world through an ideologically biased lens. What drives political polarization? It has been posited that polarization arises because holding extreme political views satisfies a need for certain and stable beliefs about the world. We examined the relationship between uncertainty tolerance and political polarization using brain-to-brain synchrony analysis, which measured committed liberals’ and conservatives’ subjective interpretation of a continuous political narrative. Participants (N=44) watched a political debate while undergoing fMRI. Shared ideology between participants increased neural synchrony in many brain areas including key regions of the valuation and theory-of-mind networks (e.g. temporoparietal junction). The degree of neural synchrony was modulated by uncertainty aversion: Uncertainty-intolerant individuals experienced greater brain-to-brain synchrony with politically like-minded peers and lower synchrony with political opponents. This effect was observed for liberals and conservatives alike. Moreover, increasing neural synchrony between committed partisans predicted subsequent polarized attitude formation about the debate after the scanning session. These results suggest that uncertainty attitudes gate the shared neural processing of political narratives, thereby fueling polarized attitude formation about hot-button issues.

## Introduction

Countries around the world are experiencing the growing strain of political polarization. Opposing partisans see the world through different eyes. Where one sees the freedom to choose, another sees murder; where one sees the right to protest, another sees violent conduct (1–3). Such a polarized perception of reality hampers bipartisan cooperation and can even undermine the basic principles of democracy (3, 4). One theory posits that polarization arises because holding extreme political views satisfies a need for certain and stable beliefs about the world (5, 6). This suggests that intolerance to uncertainty (7, 8) may play an outsized role in shaping polarized perceptions. We examine the possibility that uncertainty attitudes shape how political information is processed in the human brain: Uncertainty-intolerant individuals interpret polarizing political information through an ideologically biased ‘lens’ that produces a clear-cut perception of the issue at hand. We further test whether the neural fingerprint of these uncertainty-driven polarized perceptions—that is, increased brain-to-brain synchrony between like-minded partisans—predicts the formation of polarized attitudes.

We combine two novel techniques to measure polarized perceptions of political information. Brain-to-brain synchrony (BBS) provides a direct measure of the subjective interpretation of narrative information in groups of participants (9–11). To make this metric sensitive to more subtle differences along the ideological continuum, and to test for interactive effects between ideology and other individual differences, we combined BBS with inter-subject representational similarity analysis (IS-RSA) (12–14). This versatile approach enabled us to leverage continuous (rather than group-based) individual differences to test whether uncertainty attitudes exacerbate the ideologically biased processing of political information in the brain, and whether this fuels polarized political attitude formation.

Using a combination of targeted online and field recruiting (N=360), we invited 22 liberals and 22 conservatives to participate in a study on political cognition (Figure 1A). While undergoing fMRI, participants viewed three types of videos: a neutrally-worded news segment on a politically charged topic (abortion; PBS News), an inflammatory debate segment (police brutality and immigration; taken from the 2016 CNN Vice-Presidential debate), and a non-political video (BBC Earth; Figure 1B). Data analysis consisted of time-locking the fMRI BOLD signal to the onset of the videos and computing voxel-wise time-course correlations between each possible pairing of subjects across the entire participant pool. The resulting brain-to-brain synchrony is an established metric of the shared subjective interpretation of dynamic, naturalistic stimuli such as video narratives (10, 15, 16). We first ensured that all three videos elicited robust baseline neural synchrony between all participants, with the strongest time-course correlations occurring in sensory brain regions (Figure S1). We then analyzed variation in neural synchrony across participant dyads using IS-RSA (Figure 1D) to test three hypotheses: 1) shared ideology between subjects will predict similar neural encoding of political stimuli, 2) intolerance of uncertainty will modulate neural synchrony in committed partisans, and 3) increasing neural synchrony will predict similarity in subsequent expression of attitudes about the political stimuli.

**Figure 1.**
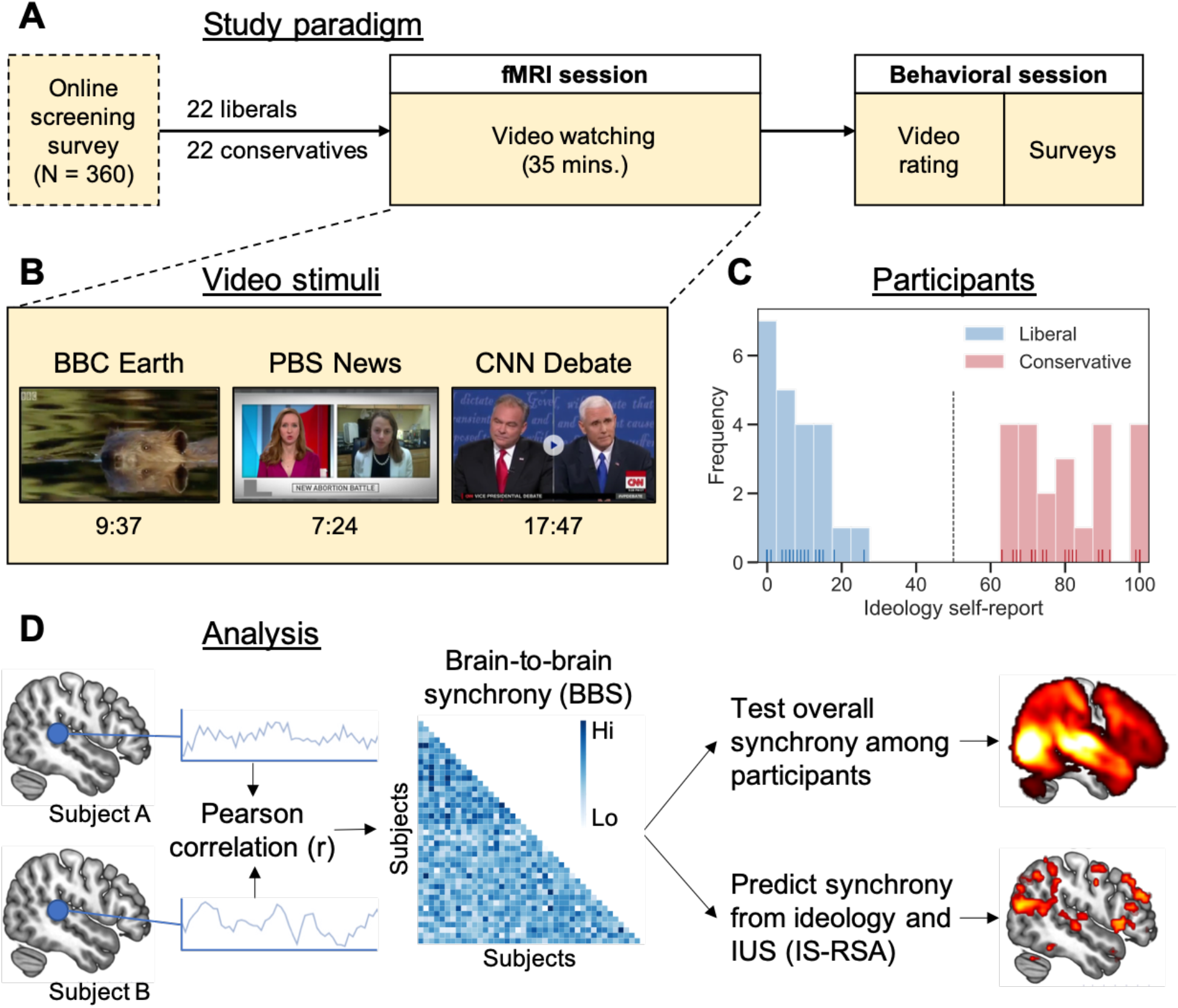
A. Participants underwent fMRI and behavioral testing as part of a larger study on political cognition. B. Participants viewed three videos in a fixed order while undergoing fMRI. C. Participants were clearly divided on political ideology. D. Analytical approach. We tested both for overall neural synchrony between participants, and for variation in neural synchrony as a function of ideology and intolerance of uncertainty (IUS). Statistical map slices taken from figures S1 and 2C. IS-RSA: inter-subject representational similarity analysis.

## Results

### Ideological similarity drives brain-to-brain synchrony

To first test whether ideological similarity drives brain-to-brain synchrony, we asked participants to self-report their political ideology using a slider (17) ranging from ‘extremely liberal’ (0) to ‘extremely conservative’ (100; Figure 1C; see Methods for validation). Inter-subject ideological closeness served as a predictor of neural synchrony in an IS-RSA model accounting for inherent statistical dependencies between pairwise observations (12). This ideological-synchrony analysis revealed no active clusters for the BBC Earth video, right angular gyrus activity for the neutrally worded PBS News abortion segment, and many clusters for the political debate video, where shared ideology was predictive of a globally synchronized brain response (Figure 2A-C). Active clusters for the debate video included regions associated with valuation (ventral striatum, medial orbitofrontal cortex) (18), theory-of-mind (temporoparietal junction, precuneus) (19), and affect (anterior insula) (20). Even though abortion is a highly polarizing topic, the neutrally-worded news video yielded much less ideology-driven neural synchrony than the inflammatory debate video, which suggests that polarized perception is not just driven by ideological differences but also by the way polarizing issues are presented.

**Figure 2.**
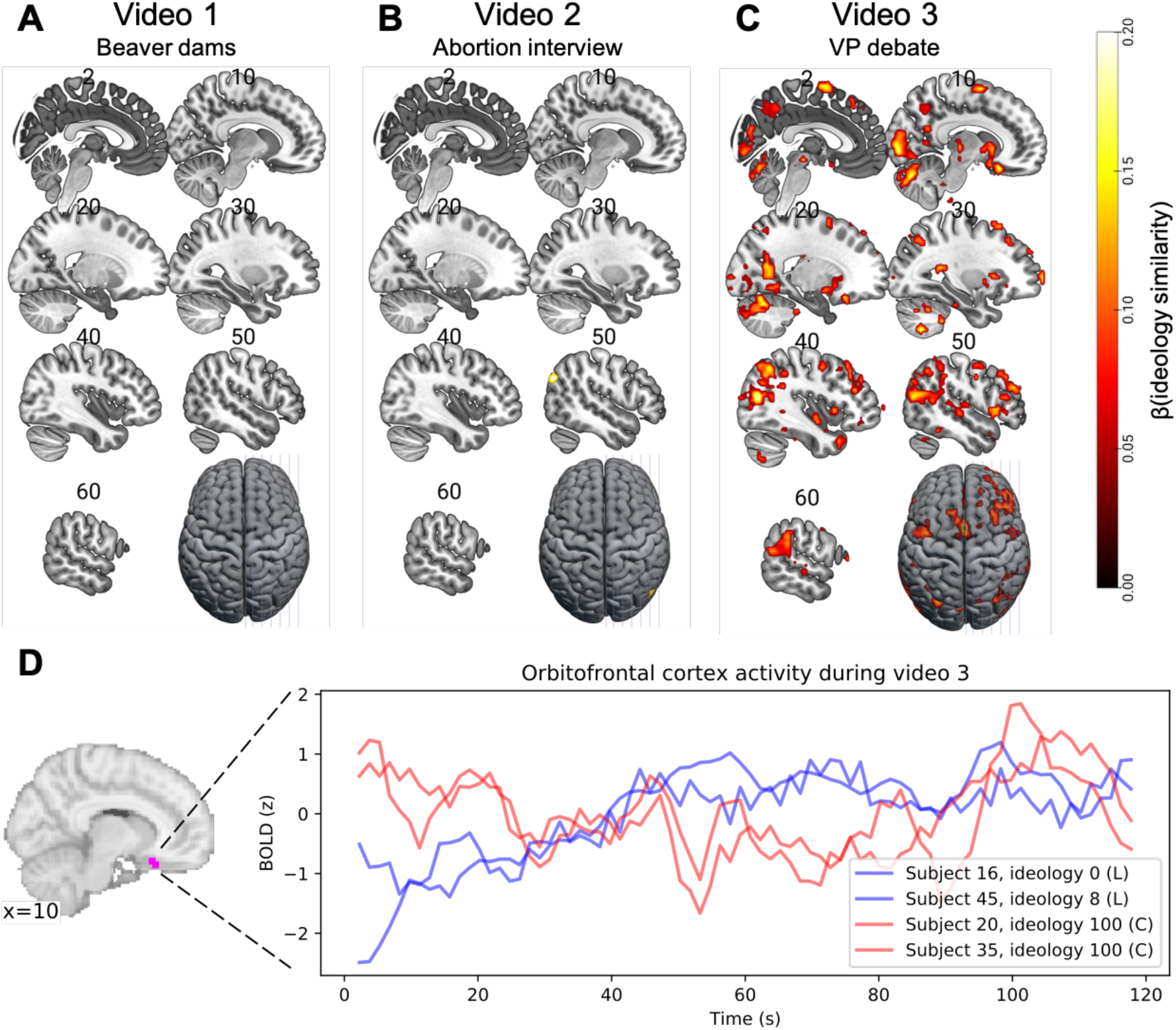
A-C. Ideology similarity drove brain-to-brain synchrony during an inflammatory political debate video. Thresholds: voxel-wise *p*(FDR) < 0.05, cluster size ≥ 5 voxels. Slice numbers indicate MNI *x* coordinate. D. Activity time-courses in medial orbitofrontal cortex were synchronized between like-minded individuals. The first two minutes of video 3 are shown, during which liberal Democrat Tim Kaine is speaking. BOLD is z-scored at the subject level; time-courses are smoothed using a 6s rolling window average to reveal trends at the intrinsic timescale of the BOLD response.

Interestingly, the synchronization of the BOLD signal was greatest between like-minded individuals regardless of whether they were both liberal or conservative, which is illustrated by representative activity time-courses of two participant dyads in the medial orbitofrontal cortex (mOFC) during the debate segment (Figure 2D). In other words, similar neural synchrony was observed in a host of brain regions across the ideological divide, revealing that sharing partisan beliefs yields similar neural encoding of a political stimulus at the time of perception.

### Intolerance of uncertainty modulates ideology-driven neural synchrony

To test whether uncertainty attitudes shape polarized information processing, we then linked neural synchrony to the well-validated 27-item intolerance of uncertainty scale (IUS) (21), which includes items like “The ambiguities in life stress me”. IUS was not associated with political ideology (ideology versus overall IUS score: *r*(42) = −0.09, *p* = 0.56), nor with ideological extremity (distance from the ideology midpoint versus overall IUS: *r*(42) = −0.06, *p* = 0.69), which makes it possible to test for independent effects of ideology and IUS on neural synchrony. We hypothesized that participant pairs with similar political views would have greater neural synchrony if they were also uncertainty-intolerant. To test this, we defined a pairwise metric of ‘joint IUS’ as the product of both IUS scores for each participant pair. We then added this as a predictor to our ideology-based IS-RSA model of neural synchrony during the debate video, positing that joint IUS would positively interact with the ideological-similarity effect.

Confirming our prediction, the ideology-IUS interaction predicted neural synchrony over and above ideological similarity in a suite of regions including bilateral temporoparietal junction (TPJ), right anterior insula (rAI), and precuneus (Figure S2C-D; Table S1). To understand the directionality of these interaction effects, we plotted the neural synchrony predicted by the fitted regression models in three of the detected regions: left TPJ, right AI, and left precuneus (Figure 3A). For each of these ROIs, intolerance to uncertainty exacerbated neural polarization, such that two individuals who were both intolerant to uncertainty and like-minded committed partisans produced significantly more neural synchrony.

**Figure 3.**
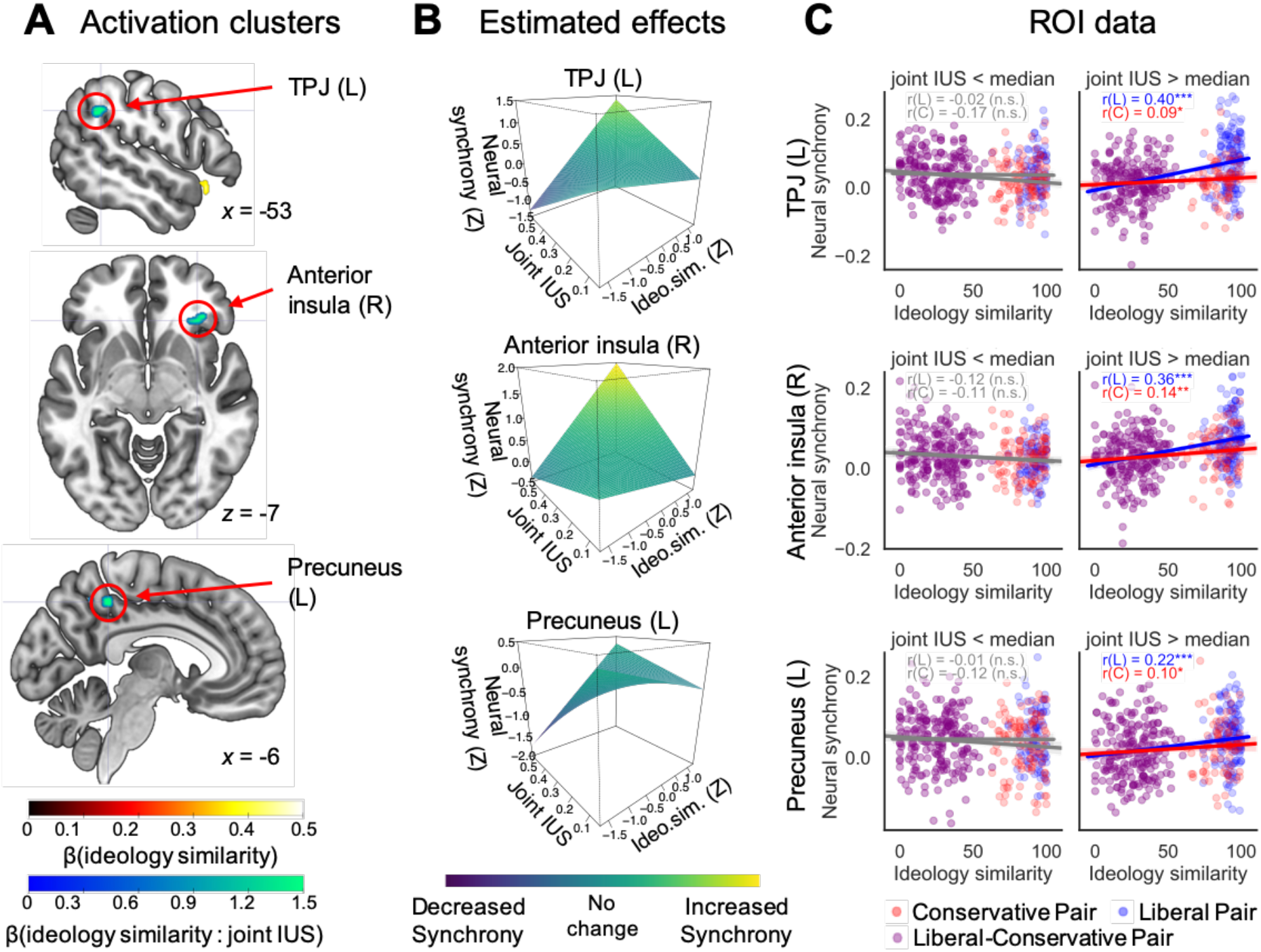
A. Neural synchrony in key socio-emotional brain regions was driven by the interaction between uncertainty-intolerance and ideology. Thresholds: voxel-wise *p*(FDR) < 0.05 and cluster size ≥ 5 voxels. B. Simulating neural synchrony from the regression model revealed that joint IUS modulated neural polarization: ideological similarity only predicted neural synchrony when participants were intolerant to uncertainty (high in joint IUS). C. Observed neural synchrony per ROI confirmed that ideological similarity only boosted synchrony in subject pairs with above-median joint intolerance of uncertainty. This effect was present for liberal (L) as well as conservative (C) ideology. *** *p* < 0.001; ** *p* < 0.01; *** *p* < 0.001 based on permutation tests. Shaded areas around lines in C. represent bootstrapped 95% confidence intervals.

To illuminate whether intolerance to uncertainty affected ideological processing writ large or was restricted to one side of the political spectrum, we tested if the IUS-modulated increase in neural synchrony was present for both liberals and conservatives. We observed that the increase in neural synchrony was driven by both ends of the political spectrum: Ideological similarity predicted increased neural synchrony for uncertainty-intolerant liberal and conservative pairs alike (Figure 3C; Supplementary Results 1). Put simply, biased information processing was not unique to any particular political persuasion, but rather influenced uncertainty-intolerant partisans across the board.

### Neural synchrony predicts post-scan polarized attitude formation

Finally, to probe whether brain-to-brain synchrony indexed the subjective interpretation of narrative information (10, 11) leading to politically relevant judgements, we examined whether synchronous brain activity between participants played an active role in polarized attitude formation about the political stimuli. We computed an inter-subject similarity score in attitude formation by measuring how much participants agreed with a set of statements made by the Vice-Presidential candidates in the debate video. We then used ideological similarity, joint IUS, their interaction, and the inter-subject neural synchrony in the three ROIs to predict attitude similarity amongst our participants. We compared this to a model without any neural data. Adding brain-to-brain synchrony significantly improved predictions of post-scan attitude expression similarity (*X^2^*(3) = 8.27, *p* = 0.041). In this model, ideological similarity positively predicted attitude similarity (*β* = 0.014 ± 0.001 (S.E.), *t*(820.1) = 18.1, one-sided *p* < 0.001), as did neural synchrony in two out of three ROIs (AI: *β* = 0.44 ± 0.18, *t*(842.8) = 2.46, *p* = 0.007; precuneus: *β* = 0.20 ± 0.14, *t*(836.2) = 1.36, *p* = 0.087; TPJ: *β* = --0.032 ± 0.15, *t*(833.1) = −0.22, *p* = 0.41). These results suggest that uncertainty-driven neural polarization plays a mechanistic role in the formation of polarized political attitudes.

## Discussion

Our findings reveal that ideological similarities are associated with increased neural synchrony between political partisans, regardless of their ideological tilt. These effects are exacerbated by increasing intolerance to uncertainty, which only enhances polarized attitude formation. Collectively, this provides neural evidence that uncertainty-intolerant individuals are more susceptible to holding rigid political beliefs (5–7), and that this uncertainty fuels how politicized information becomes polarized. In our experiment, uncertainty-intolerant individuals experienced greater brain-to-brain synchrony with politically like-minded peers, regardless of whether they were liberal or conservative, and lower synchrony with political opponents. This suggests that uncertainty-intolerant individuals see the political world through a stronger partisan lens, construing a more biased picture of the political reality (4, 22). Rather than ideology alone, cognitive traits such as intolerance to uncertainty—which interact with ideology to form a polarized perception of the world—may be the lynch pin of political polarization.

This work extends recent research on neural polarization (9) in several important ways. Our data indicates that neural polarization may only arise when political information is presented in a polarizing way (e.g. a debate as opposed to a neutrally-worded political news item). Moreover, we show that the effect of ideology on brain-to-brain synchrony is largely driven by participants who are highly averse to uncertainty. This implies that polarized perception is not irremediable but depends on additional factors that vary between stimuli, contexts, and individuals. The growing uncertainty caused by large-scale societal events in the past year (e.g. job loss and a global pandemic) may fuel political polarization by sowing rigidly partisan perceptions of the world. Conversely, interventions against polarization may be successful by addressing citizens’ sources of worry (8, 23).

## Methods

### Participants

The data analyzed for this paper were collected as part of a larger study on political preferences. For this study, 360 potential participants completed a screening survey prompted by online advertisements, paper flyers, personal visits to political meetings throughout the state, or word-of-mouth. The slider measure of ideology (17) was administered in the screening survey. Based on this measure, we invited 22 self-reported conservatives and 22 liberals for an in-lab session, all of whom were right-handed and eligible for MRI. One participant was excluded from the analysis because they indicated a different ideology on the screening survey versus the post-scan political survey battery (see below), leaving 21 conservatives (13 men and 8 women; age range 18-61, mean 36 y ± s.d. 15 y) and 22 liberals (13 men and 9 women; age range 18 to 60, mean age 28 ± 12 y) representing a range of ideological extremity (see Figure 1C). Age nor gender differed significantly between the two groups (all *p*s > 0.05), but to ensure that our effects were not driven by demographic differences we controlled for them in several control analyses (Supplementary Figures 3 and 4). The two groups were matched on education level (number of years completed) and annual income (two-sample t-tests, two-sided: all *p*s > 0.4). All participants provided written informed consent and were paid for their participation in the study. The study procedures were approved by the local ethics committee.

### Procedure

Participants came to the lab for one session of about 3 hours. A subset of the tasks completed in this time were analyzed for the current paper. The first half of the study session took place in the functional MRI laboratory. Upon arrival, participants read instructions about the experiment, answered comprehension questions, and confirmed or updated their MRI screening information. Participants then entered the MRI scanner for a scanning session of about 1.5 hours, with soft padding positioned around their head to minimize head motion. In the scanner, participants completed a cognitive task (not analyzed here), underwent a 5-minute anatomical scan, and completed the video watching task described in the Main Text. During the scans, participants wore two electrodes on their non-dominant (left) hand to measure skin conductance. They held a button response box in their right hand. In between each scanner run or task block, the experimenter verbally communicated with the participants to ensure that they were comfortable and attentive. The video watching task consisted of a fixed sequence of three videos, which can be viewed at the following web pages: a) BBC Earth, “Beaver Lodge Construction Squad”: https://www.youtube.com/watch?v=iyNA62FrKCE; b) PBS News Hour, “ State battles over abortion policy anticipate a post-Roe world”: https://www.pbs.org/newshour/show/state-battles-over-abortion-policy-anticipate-a-post-roe-world; c) CNN, “Vice-Presidential Debate 2016” between liberal Democrat Tim Kaine and conservative Republican Mike Pence, clip from about 24:35 to 42:10: https://www.youtube.com/watch?v=ox8PTXwDYdc.

After participants were taken out of the scanner, the experimenter walked with them to a different building on campus for behavioral testing. During this session, participants first completed a survey of their comprehension and judgment of the videos. Judgment items measured attitudes about statements made in the debate video, such as Mike Pence’s statement *the immigration reform plan of Hillary Clinton and Tim Kaine amounts to “amnesty”*, with a seven-point Likert-type response scale ranging from “Strongly disagree” to “Strongly agree”. Next, participants completed several cognitive tasks (not analyzed here) and an extensive survey battery of 5 political and 3 cognitive questionnaires. The political questionnaires were the updated social and economic conservatism scale (24) (SECS), the Schwartz short values survey (25) (SSVS), right-(26) and left-wing(27) authoritarianism surveys (RWA and LWA, respectively), and a short-form social dominance orientation survey (28) (SDO). A principal component analysis (PCA) on these five political survey measures revealed a first component that accounted for 37% of variance across all survey items and was very strongly correlated to the ideology scale measure from the screening survey (r(41) = 0.89, p < 0.001). This strong relationship across time and surveys validates the ideology scale used to recruit participants as a reliable metric of fundamental political orientation. The PCA also identified one participant as a strong outlier, as he had rated his ideology as strongly conservative on the screening slider measure but scored more than three standard deviations below the mean of the conservative group on the first component of the PCA (in fact scoring squarely among the liberals); this participant was therefore excluded from analysis. The cognitive questionnaires included the intolerance of uncertainty scale (21) (IUS; see Main Text), a short need for closure scale (29) (NFC), and the interpersonal reactivity index (30) (IRI). IUS item scores were averaged for each participant and normalized to a 0-1 range. Just as individual-level IUS was not associated with ideology (see Main Text), pairwise ideology similarity was not associated with joint IUS (*r*(859) = 0.035, *p* = 0.30) nor with pairwise cosine similarity in responses across the entire intolerance of uncertainty survey (*r*(859) = 0.023, *p* = 0.50). NFC and IRI were not analyzed for this paper.

### fMRI acquisition

MR images were collected on a Siemens Prisma Fit 3-Tesla research-dedicated scanner. T2*-weighted functional scans were acquired using a multi-slice sequence capturing three slices at once to ensure whole-brain coverage with short repetition time (TR = 1500 ms), which increases the number of time points and thus statistical power for brain-to-brain temporal synchrony analysis. 60 3-mm transverse slices were acquired, each with 64×64 voxels of 3.0 mm isotropic, building up a field of view (FOV) that covered the entire brain except part of the cerebellum. The FOV was tilted upward by 25 degrees at the front of the brain to minimize tissue gradient-related signal dropout in the orbitofrontal cortex. Contrast settings were optimized for cortical grey matter (TE = 30 ms, flip angle = 86°). T1-weighted anatomical scans were acquired using a standard MPRAGE sequence (160 sagittal slices with 256×256 voxels of 1.0 mm isotropic, TR = 1900 ms, TE = 3.02 ms, flip angle = 9°).

### fMRI preprocessing

Results included in this manuscript come from preprocessing performed using *fMRIPrep* 1.5.1rc2 (31, 32) (RRID:SCR_016216), which is based on *Nipype* 1.3.0-rc1 (33, 34) (RRID:SCR_002502).

### Anatomical data preprocessing

The T1-weighted (T1w) image was corrected for intensity non-uniformity (INU) with N4BiasFieldCorrection (35), distributed with ANTs 2.2.0(36) (RRID:SCR_004757), and used as T1w-reference throughout the workflow. The T1w-reference was then skull-stripped with a *Nipype* implementation of the antsBrainExtraction.sh workflow (from ANTs), using OASIS30ANTs as target template. Brain tissue segmentation of cerebrospinal fluid (CSF), white-matter (WM) and gray-matter (GM) was performed on the brain-extracted T1w using fast (37) (FSL 5.0.9, RRID:SCR_002823). Volume-based spatial normalization to one standard space (MNI152NLin2009cAsym) was performed through nonlinear registration with antsRegistration (ANTs 2.2.0), using brain-extracted versions of both T1w reference and the T1w template. The following template was selected for spatial normalization: *ICBM 152 Nonlinear Asymmetrical template version 2009c* (38) [RRID:SCR_008796; TemplateFlow ID: MNI152NLin2009cAsym].

### Functional data preprocessing

For each of the 9 BOLD runs found per subject (across all tasks and sessions), the following preprocessing was performed. First, a reference volume and its skull-stripped version were generated using a custom methodology of *fMRIPrep*. The BOLD reference was then co-registered to the T1w reference using flirt (39) (FSL 5.0.9) with the boundary-based registration (40) cost-function. Co-registration was configured with nine degrees of freedom to account for distortions remaining in the BOLD reference. Head-motion parameters with respect to the BOLD reference (transformation matrices, and six corresponding rotation and translation parameters) are estimated before any spatiotemporal filtering using mcflirt (41) (FSL 5.0.9). BOLD runs were slice-time corrected using 3dTshift from AFNI 20160207 (42) (RRID:SCR_005927). The BOLD time-series (including slice-timing correction when applied) were resampled onto their original, native space by applying a single, composite transform to correct for head-motion and susceptibility distortions. These resampled BOLD time-series will be referred to as *preprocessed BOLD in original space*, or just *preprocessed BOLD*. The BOLD time-series were resampled into standard space, generating a *preprocessed BOLD run in MNI152NLin2009cAsym space*. First, a reference volume and its skull-stripped version were generated using a custom methodology of *fMRIPrep*. Several confounding time-series were calculated based on the *preprocessed BOLD*: framewise displacement (FD), DVARS and three region-wise global signals. FD and DVARS are calculated for each functional run, both using their implementations in *Nipype* (following the definitions by Power et al. (43)). The three global signals are extracted within the CSF, the WM, and the whole-brain masks. Additionally, a set of physiological regressors were extracted to allow for component-based noise correction (*CompCor* (44)). Principal components are estimated after high-pass filtering the preprocessed BOLD time-series (using a discrete cosine filter with 128s cut-off) for the two *CompCor* variants: temporal (tCompCor) and anatomical (aCompCor). tCompCor components are then calculated from the top 5% variable voxels within a mask covering the subcortical regions. This subcortical mask is obtained by heavily eroding the brain mask, which ensures it does not include cortical GM regions. For aCompCor, components are calculated within the intersection of the aforementioned mask and the union of CSF and WM masks calculated in T1w space, after their projection to the native space of each functional run (using the inverse BOLD-to-T1w transformation). Components are also calculated separately within the WM and CSF masks. For each CompCor decomposition, the k components with the largest singular values are retained, such that the retained components’ time series are sufficient to explain 50 percent of variance across the nuisance mask (CSF, WM, combined, or temporal). The remaining components are dropped from consideration. The head-motion estimates calculated in the correction step were also placed within the corresponding confounds file. The confound time series derived from head motion estimates and global signals were expanded with the inclusion of temporal derivatives and quadratic terms for each (45). Frames that exceeded a threshold of 1.0 mm FD or 100.0 standardised DVARS were annotated as motion outliers. All resamplings can be performed with a single interpolation step by composing all the pertinent transformations (i.e. head-motion transform matrices, susceptibility distortion correction when available, and co-registrations to anatomical and output spaces). Gridded (volumetric) resamplings were performed using antsApplyTransforms (ANTs), configured with Lanczos interpolation to minimize the smoothing effects of other kernels (46). Non-gridded (surface) resamplings were performed using mri_vol2surf (FreeSurfer).

### Video fMRI data cleaning

For video 1, two participants’ data were excluded from analysis, one due to falling asleep in the scanner and one due to excessive head motion. For video 3, one participants’ data were excluded from analysis due to excessive head motion. All other functional fMRI data was further preprocessed using *nltools* 0.3.14 (47) to remove signal components related to motion and other sources of noise. To this end, general linear models of each voxel’s signal time series were constructed with the following regressors: average CSF signal; average white matter signal; the six realignment parameters, their derivatives, their squares, and their squared derivatives (48); zero-, first-, and second-order polynomials for removal of intercepts and linear/quadratic trends; and a regressor for each motion spike, which has value 1 at the TR where the spike was detected (FD > 1 mm) and zeros elsewhere. The residual time series of each voxel were then used for statistical analysis.

### Statistical fMRI analysis

We first established that all three videos elicited robust baseline neural synchrony between all participants (known as inter-subject correlation; ISC (11)). To this end, we correlated each participant’s signal time course in a voxel to the average time course of all other participants. The resulting distribution of *r* values was subjected to a sign-flip permutation test (5000 permutations) to see whether its average value was greater than would be expected by chance (9, 11). Voxel-wise p-values were thresholded at a false-discovery rate of 0.001. ISC effects were significant throughout the brain, with the strongest time-course correlations occurring in sensory brain regions (Figure S1).

For the inter-subject representational similarity analyses (IS-RSA) (14), we wrote a custom implementation of the mixed-effects regression approach reported by Chen et al. (12) based on the packages *lme4* 1.1-23 and *lmerTest* 3.1-2 for R 3.5.2. In this analysis, we first computed the temporal synchrony of the BOLD signal time courses (Pearson correlation) between each pair of subjects for each voxel. At each voxel, this vector of pairwise synchrony values was then regressed onto a set of regressors that includes one or more fixed effects (for instance ‘ideology similarity’ or ‘joint IUS’) and random participant intercepts. Since each observation in the mixed-effects regression corresponded to a unique *pair* of subjects, the model for each observation includes a random participant intercept for *both* participants involved in that participant pair (12). For the first IS-RSA, we regressed brain-to-brain synchrony onto ideology similarity, which was computed as 100 − *abs*(*ideology*_1_ − *ideology*_2_) where 1 and 2 refer to the two participants in the current participant pair and z-scored. The signal time courses in Figure 2D were generated from the mean BOLD time series in a spherical ROI with radius = 6 mm centered around the peak voxel for the ideology effect in OFC. For the second IS-RSA, we regressed neural synchrony onto ideology similarity and joint IUS, the latter of which is the product of both participants’ total intolerance of uncertainty scores normalized to a 0-1 range. By also including the interaction effect between these two regressors, we could test whether IUS modulated the effect of ideology on brain response. Regressor beta maps were thresholded at false-discovery rate (FDR)-corrected *p*-values of 0.05 using the *p.adjust* function in R. Surviving clusters were reported if they contained five or more contiguous voxels.

For the region-of-interest analyses in Figure 3B and 3C, we first extracted mean activity time-courses in each ROI for each participant and recomputed the brain-to-brain synchrony in these mean signals (’ROI-level neural synchrony’). As ROIs, we used activation clusters from the whole-brain ideology-IUS IS-RSA analysis. The precuneus and temporoparietal junction ROIs corresponded to our preregistered regions of interest ([URL]) and were previously found to process subjective interpretations of narrative content (10); the anterior insula ROI fits with what we know about intergroup social influence on emotion processing (49). For the estimated effect plots in Figure 3B, we re-ran the inter-subject representational similarity analysis with ideology, IUS, and their interaction on ROI-level neural synchrony. We then used the effect estimates from the fitted mixed-effects regression models to simulate the DV (neural synchrony) at a dense grid of ideology-similarity and joint-IUS values, whose axis limits matched those observed in the dataset. For the scatter plots and correlation analyses in Figure 3C, we labeled all participant dyads as containing two conservatives (CC), two liberals (LL), or one conservative and one liberal (CL). To evaluate whether the ideology-IUS interaction effect on neural synchrony was present for both conservatives and liberals, we then tested whether there was a significant correlation between ideology and neural synchrony in high-IUS pairs (joint IUS > median), a) when leaving out LL pairs and b) when leaving out CC pairs. We established significance by re-running these correlations after randomly permuting the data labels 10,000 times and comparing the true correlation coefficient to the resulting null distribution. A reanalysis that treats joint IUS as continuous rather than dichotomous yielded the same result (Supplementary Note 1).

To test whether neural activity in the obtained activation clusters was predictive of attitude formation, we added ROI-level neural synchrony to a dyadic regression model of attitude formation, where the dependent variable is the inter-subject agreement on six statements made by Tim Kaine and Mike Pence during video 3 (e.g. “Law enforcement in this country is a force for good”). Inter-subject agreement was computed as the cosine similarity between the 6-item response vectors of each pair of participants. This variable was then regressed on ideology similarity, joint IUS, and their interaction effect, as well as the regular random subject intercepts (model 1). We additionally regressed it onto a model that included the same regressors as model 1, but also ROI-level neural synchrony in each of the three ROIs (model 2). We compared the explained variance of model 2 versus model 1 using the *anova* function in R.

### Control analyses

It is well-established that similarities on demographic and social variables can increase inter-subject synchrony in brain responses to video stimuli (50). Although this potential confound cannot account for the ideology-IUS interaction effects that were observed in our key analyses, we nevertheless wanted to ensure that the effects reported here were specific to political polarization. To this end, we re-ran the inter-subject RSA analyses with additional regressors, each controlling for a demographic or experimental factor that may impact neural synchrony: age, gender, undergraduate student status, sampling source (from the university or from the community), and scan day (participants were scanned across a ~6-month period). For the ideology model, no clusters survived this additional control for video 2, but nearly all of the many active clusters survived for video 3 (Figure S3). For the ideology-IUS interaction model, all clusters survived except the precuneus (Figure S4).

## Data availability

All data needed to reproduce the results in this paper will be made available on a public repository upon publication.

## Code availability

All analysis code needed to reproduce the results in this paper will be made available on a public repository upon publication.

## Acknowledgements

This work was supported by Brown University OVPR Research Seed Fund Award GR300152 and NIH COBRE grant P20GM103645. We thank Uri Hasson for fruitful discussions during the study design and analysis pipeline. We are grateful to Pedro Rodriguez, Daantje de Bruin, Rose McDermott and the FeldmanHall Lab for productive discussions on study design and interpretation. Finally, the contributions of Wenning Deng, Marlon Sherman, Rachel Yan, Logan Bickel, Avinash Vaidya, Romy Froemer, Amrita Lamba, Lynn Fanella, and Fabienne McEleney were invaluable during data collection.

